# Automated high-quality reconstruction of metabolic networks from high-throughput data

**DOI:** 10.1101/282251

**Authors:** Daniel Hartleb, C. Jonathan Fritzemeier, Martin J. Lercher

## Abstract

While new genomes are sequenced at ever increasing rates, their phenotypic analysis remains a major bottleneck of biomedical research. The generation of genome-scale metabolic models capable of accurate phenotypic predictions is a labor-intensive endeavor; accordingly, such models are available for only a small percentage of sequenced species. The standard metabolic reconstruction process starts from a (semi-)automatically generated draft model, which is then refined through extensive manual curation. Here, we present a novel strategy suitable for full automation, which exploits high-throughput gene knockout or nutritional growth data. We test this strategy by reconstructing accurate genome-scale metabolic models for three strains of *Streptococcus*, a major human pathogen. The resulting models contain a lower proportion of reactions unsupported by genomic evidence than the most widely used *E. coli* model, but reach the same accuracy in terms of knockout prediction. We confirm the models’ predictive power by analyzing experimental data for auxotrophy, additional nutritional environments, and double gene knockouts, and we generate a list of potential drug targets. Our results demonstrate the feasibility of reconstructing high-quality genome-scale metabolic models from high-throughput data, a strategy that promises to massively accelerate the exploration of metabolic phenotypes.

**Significance statement:** Reading bacterial genomes has become a cheap, standard laboratory procedure. A genome by itself, however, is of little information value – we need a way to translate its abstract letter sequence into a model that describes the capabilities of its carrier. Until now, this endeavor required months of manual work by experts. Here, we show how this process can be automated by utilizing high-throughput experimental data. We use our novel strategy to generate highly accurate metabolic models for three strains of *Streptococcus*, a major threat to human health.

## Introduction

Genome-scale metabolic models have been reconstructed for a wide range of organisms (1); they have been used successfully to predict metabolic phenotypes of prokaryotic and eukaryotic cells, in applications ranging from evolutionary studies (2-4) to bioengineering designs (5). An initial draft reconstruction is typically created by mapping each gene of the organism of interest to a closely related, well-annotated relative or to a database of known metabolic functions (6), a task that can easily be automated. These draft models are often unable to produce biomass. Gap-filling methods are applied to ensure functionality of the networks, adding reactions without genomic evidence (7).

The currently most widely applied methodology to generate a reliable genome-scale metabolic network can be considered a hybrid strategy: it starts from an automatically generated draft model based on sequence similarities, followed by manual refinement of reaction content and gene-reaction mapping (6). This mode of metabolic network reconstruction is still time-consuming and laborious. To accelerate the development of metabolic models, automatic and semi-automatic algorithms have been developed (8-10). However, the resulting metabolic networks are typically unreliable and require extensive manual curation (6). For example, reactions may be missing or were added without evidence to facilitate biomass formation. Furthermore, reactions may erroneously be assumed reversible or may have been assigned to the wrong compartment, gene-reaction associations may be incorrect, or biomass components may be missing (11). Accordingly, growth/non-growth predictions for gene knockout strains from automated reconstructions typically show Matthews correlation coefficients *R*_*M*_<0.5, while extensively manually curated models often agree much better with experimental data (e.g., *R*_*M*_=0.67 for the iJO1366 *E. coli* model (12), where *R*_*M*_ =1 wouldcorrespond to a perfect prediction).

Automatic reconstructions suffer from being based on only one or a few nutritional environments in which the organism is supposed to grow. They lack the capability to utilize – equally important – information about environments in which the investigated organism cannot grow. Moreover, automatic reconstruction tools possess no mechanism to automatically incorporate genome-wide knock-out data, although such datasets are usually highly informative and thus might lead to more accurate automatic reconstructions.

Here, we develop a pipeline for the automated reconstruction of high-quality genome-scale metabolic networks. We submit an automatically generated draft network to a bi-level optimization algorithm that minimizes the deviation between model predictions and experimental growth/non-growth data from sets of gene knockout strains and/or nutritional environments. We apply our method to three species of *Streptococcus*, gram-positive bacteria that pose a sever risk to human health. *Streptococcus pyogenes* is a Lancefield group A streptococcus (GAS) (13), and is one of the two clinically most important human pathogens (14). *S. pyogenes* are responsible for at least 616 million cases of throat infection (pharyngitis, tonsillitis) worldwide per year, and 111 million cases of skin infection (primarily non-bullous impetigo) in children of less developed countries (15). *Streptococcus sanguinis* (formerly known as *S. sanguis*) is categorized as Lancefield group H (16). These oral bacteria can enter the bloodstream and may cause severe endocarditis (17, 18). Finally, *Streptococcus agalactiae* is a Lancefield group B bacterium, and is one of the major causes of pneumonia and meningitis in neonates (19, 20).

## Results

### Automated high-quality metabolic reconstruction strategy

Analogous to the hybrid strategy for metabolic network reconstruction currently widely employed, we start with the automated construction of a draft model based on sequence similarity. We do this by successively submitting the gene sequences of the organism under study to different annotation sources that associate them with metabolic functions (Figure 1):(i) KBase (which implements the functionality of Model SEED (8)); (ii) the existing metabolic model of *Lactococcus lactis* (21), a close relative of Streptococci; (iii) TransportDB (22); and (iv) KEGG/KAAS (23). We ordered these information sources by increasing reliability; accordingly, at each step, information on metabolic gene functions from previous steps is superseded by newer information.

**Figure 1.**
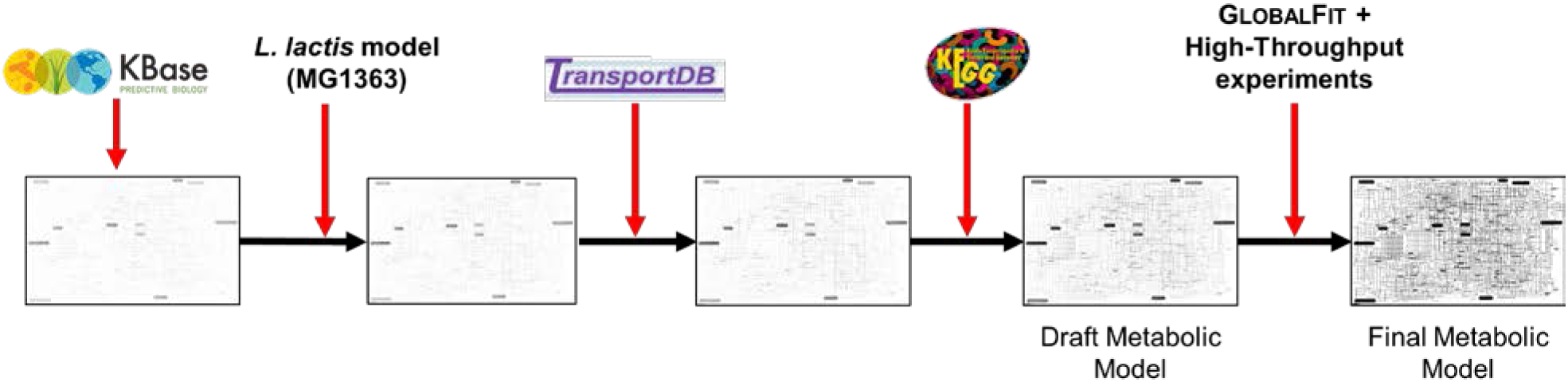
Automated workflow used for the reconstruction of the *Streptococcus* metabolic models. Information added at each step supersedes information from previous steps.

In contrast to the standard hybrid automatic/manual strategy, we continue with an automated refinement of the resulting draft model, informed through comparisons to viability data for gene knockouts and/or nutritional environments (Figure 1). For the application to *Streptococcus*, we only used gene knockout data. We identified false viability predictions of the draft model for individual gene knockouts using flux balance analysis (FBA) (24) with the biomass reaction provided by the *L. lactis* metabolic model. FPp (false positive predictions) are cases where the *in silico* gene knockout simulation predicted growth, while the *in vivo* experiment showed no growth. FNp (false negative predictions) are cases where the *in silico* analysis predicted no growth, while the corresponding knockout strain was viable in the experiment.

The automated refinement was carried out using the GLOBALFIT algorithm, which was originally developed for the further reconciliation of high-quality metabolic network reconstructions with experimental viability data (12). GLOBALFIT is a bi-level optimization program that identifies smallest sets of network modifications in order to minimize the number of FNp and FPp cases; allowed network modifications are (i) removals or (ii) reversibility changes of existing reactions;(iii) additions of reactions to the model from a database of potential reactions; (iv) removals of metabolites from the biomass; and (v) additions of metabolites to the biomass.

The strength of GLOBALFIT in comparison to previous approaches is its ability to consider several growth and/or non-growth cases simultaneously. In particular, when considering FPp, it is important to simultaneously consider a true growth case to avoid trivial solutions such as the removal of an essential reaction (12).

### Generation of high-quality metabolic models for streptococci

To build the *Streptococcus* metabolic reconstructions, we initially solved each FPp simultaneously with a wild type growth case and each FNp simultaneously with a non-growth case. If the network changes suggested by GLOBALFIT introduced new errors by converting true positive predictions (TPp) to FNp, or true negative predictions (TNp) to FPp, we solved the examined case again, this time simultaneously with all distorted cases.

To create a conservative set of potential additional enzymatic reactions, we first inferred homologs between each *Streptococcus* genome and the genes included in metabolic reconstructions in the BiGG database (25). Genes were considered homologous if bi-directional BLAST searches associated the genes with e-values <10^−13^. We only allowed the addition of a reaction to the *Streptococcus* model if homologs to genes sufficient to catalyse the reaction in one of the BiGG models were present in the *Streptococcus* genome.

We allowed the potential reversal of irreversible reactions only for those reactions that were classified at least as “reversible with uncertainty” in the *E. coli* metabolic network (26). Biomass reaction changes (addition or removal of biomass components) were only introduced if no other model changes could rescue the FNp or FPp. The quantitative contribution of each biomass component to the total biomass will have to be manually adjusted to allow quantitative predictions of biomass yield, especially for biomass components added during the network refinement with GLOBALFIT.

The resulting metabolic network for *S. pyogenes* contains 653 metabolites and 661 reactions, accounting for 455 genes. The predicted network for *S. sanguinis* is substantially larger, containing 805 metabolites and 840 reactions, corresponding to 597 genes. The *S. agalactiae* model encompasses 653 metabolites and 661 reactions, accounting for 455 genes. Because streptococci are gram positive bacteria, each genome-scale metabolic network contains only two compartments: extracellular and cytosolic.

The largest set of FPp in all networks is due to the experimentally observed essentiality of the F-ATPase complex, which is encoded by 8 genes and is part of the respiratory chain of many organisms. *S. sanguinis, S. pyogenes*, and *S. agalactiae* lack a respiratory chain, and do not use this enzyme to produce ATP; instead, the F-ATPase consumes ATP to pump protons out of the cell to maintain an internal pH that is more basic than the exterior (27). Thus, the F-ATPase does not have any metabolic function that can be accounted for in FBA models, explaining why even a perfect FBA representation of *Streptococcus* metabolism cannot predict the essentiality of the corresponding genes.

The networks predict viability of gene knockouts with high accuracy (Table 1). On average, we obtain an accuracy (percentage of true predictions) of 94.8%. Matthew’s correlation coefficient (28), a more balanced measure of prediction quality, is on average *R*_*M*_=0.85. When discounting the genes involved in the F-ATPase complex (which cannot be predicted correctly in any FBA model), average accuracy increases to 96.2%, with *R*_*M*_=0.89.

**Table 1.**
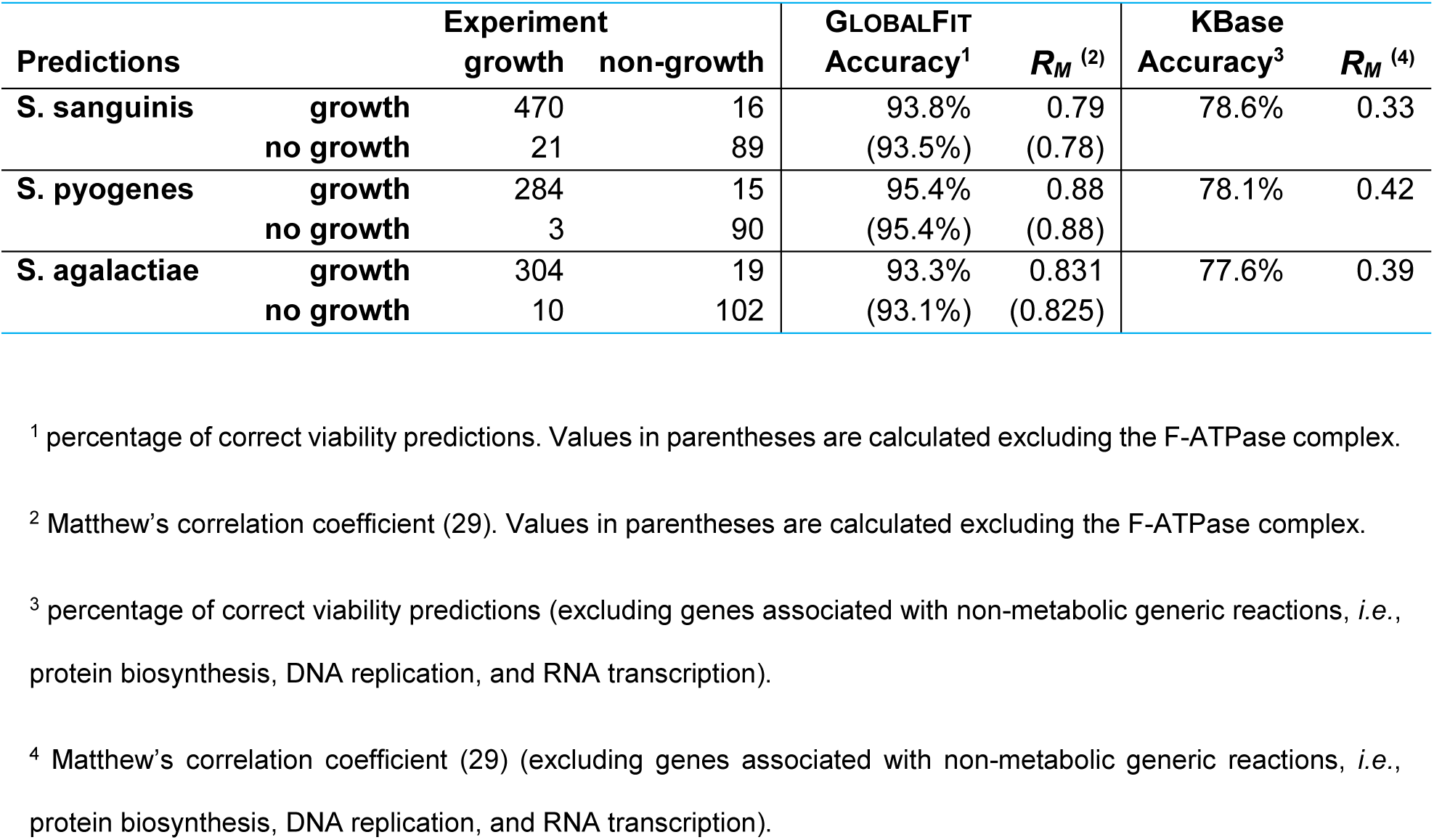
Comparison of experimental and predicted viability for Streptococci gene knockouts

| Predictions |  | Experiment |  | GLOBALFIT |  | KBase |  |
| --- | --- | --- | --- | --- | --- | --- | --- |
| | | growth | non-growth | Accuracy<sup>1</sup> | $R_M^{(2)}$ | Accuracy<sup>3</sup> | $R_M^{(4)}$ |
| **S. sanguinis** | **growth** | 470 | 16 | 93.8% | 0.79 | 78.6% | 0.33 |
|  | **no growth** | 21 | 89 | (93.5%) | (0.78) |  |  |
| **S. pyogenes** | **growth** | 284 | 15 | 95.4% | 0.88 | 78.1% | 0.42 |
|  | **no growth** | 3 | 90 | (95.4%) | (0.88) |  |  |
| **S. agalactiae** | **growth** | 304 | 19 | 93.3% | 0.831 | 77.6% | 0.39 |
|  | **no growth** | 10 | 102 | (93.1%) | (0.825) |  |  |

<sup>1</sup> percentage of correct viability predictions. Values in parentheses are calculated excluding the F-ATPase complex.
<sup>2</sup> Matthew's correlation coefficient (29). Values in parentheses are calculated excluding the F-ATPase complex.
<sup>3</sup> percentage of correct viability predictions (excluding genes associated with non-metabolic generic reactions, *i.e.*, protein biosynthesis, DNA replication, and RNA transcription).
<sup>4</sup> Matthew's correlation coefficient (29) (excluding genes associated with non-metabolic generic reactions, *i.e.*, protein biosynthesis, DNA replication, and RNA transcription).

Currently three metabolic models of *Streptococcus* strains are available (29, 30), but only the models from the AGORA project were obtainable in a usable format; the third model is only available as a PowerPoint file. The AGORA models were automatically reconstructed from the human gut microbiome. These two published models for *S. sanguinis and S. agalactiae* were tested for the same gene knockouts, but had lower accuracies and Matthew’s correlation coefficients (see Supplemental Table S7).

### Accuracy of the automatically generated Streptococcus models

To further investigate the metabolic capabilities of the three strains, we used GLOBALFIT to predict a minimal medium. The minimal medium for *S. sanguinis* contains nine nutrients, while *S. pyogenes* and *S. agalactiae* require a minimum of 22 metabolites, mainly because the latter two species require a larger number of externally supplied amino acids. To explore this issue further, we used GLOBALFIT to predict all amino acids that cannot be produced from a minimal medium from which all amino acids were removed; for *S. pyogenes* and *S. sanguinis*, this reduced minimal medium consisted of the eight nutrients glucose, phosphor, sulphur, iron, pyridoxal, niacin, riboflavin, and pantothenate. *S. agalactiae* additionally required thiamine. Our simulations on these reduced minimal media showed that *S. sanguinis* is only auxotrophic for Cysteine, while *S. pyogenes* and *S. agalactiae* are auxotrophic for 13 and 11 amino acids, respectively; in addition, *S. agalactiae* requires at least two out of four further amino acids (See Supplemental Table S1). In comparison, *L. lactis* is auxotrophic for Leucine, Histidine, and Methionine (21). The two previously published *Streptococcus* models (29) (AGORA models in the following) are only auxotrophic for three and 11 amino acids for *S. sanguinis and S. agalactiae, respectively* (see Supplemental Table S1). This table also shows observations from previous experimental studies (31-33). The new S. *sanguinis* model predicts the auxotrophy for Cysteine correctly, while auxotrophies for Isoleucine, Leucine and Valine are missing. Here predicts the AGORA model additional auxotropies for Asparagine and Methionine. The new S. *pyogenes* model predicts three out of four (non)-auxotrophies correctly. The S. *agalactiae* are consistent with the AGORA model and the experiments exept for the predicted double auxotrophies, which were not tested experimentally, and the Glutamate auxotrophy (see Supplemental Table S1 for the complete data).

We further tested the *S. sanguinis* model by comparing its predictions to growth experiments on different defined media. The metabolic model constructed by GLOBALFIT successfully predicted growth on B 48 (34) as well as on SY and M3 media (35). The study of Rogers also provided growth information on a set of SY and M3 media lacking one metabolite (either riboflavin, pantheonate, thiamine, nicotinic acid, pyridoxal, folic acid, aminobenzoic acid, or biotin). We successfully predicted the essentiality and non-essentiality of these metabolites, except for the essentiality of pyridoxal and riboflavin. We added these components to the biomass objective function; if the growth environments had been included in the original GLOBALFIT run, this addition would have occurred automatically. Pyridoxal and riboflavin were also part of the biomass reaction of the alternative network reconstructed by KBase (see below).

In a previous study, knockout mutants for three *S. sanguinis* metabolic enzymes were unexpectedly found to be viable (36). For each of these genes, Xu *et al.* identified isozymes or paralogs in the *S. sanguinis* genome. The corresponding double-gene knockouts were unviable for two of the three reactions, but were unexpectedly viable for one enzyme (NAD(P)H-dependent glycerol-3-phosphate dehydrogenase). While we did not include these double knockouts in the training set used to derive the *S. sanguinis* network, all three double knockouts were correctly predicted by our model.

To benchmark the predictive power of our reconstructed metabolic models, we compared them to the network automatically reconstructed from the genome sequence by KBase. The KBase model for *S. agalactiae* was not able to grow anaerobically and was incapable of producing NADP; thus, we allowed the influx of oxygen and NADP for the corresponding simulations. All three networks were substantially less accurate in predicting gene knock-outs (accuracy ≤79%, R*M* ≤0.42 for metabolic reactions; Table 1 and Supplemental Table S2). The F-ATP-synthase complex was not included in any metabolic network by KBase.

Could the superior performance of the GLOBALFIT models be due to overfitting to the gene knockout data? To test this, we also compared amino acid auxotrophy (33-35, 37), which was not used by either network reconstruction approach. While GLOBALFIT predictions were fully consistent with the experimental results, the KBase model required more amino acids than experimentally observed (Supplemental Table S3). For example, KBase predicted *S. sanguinis* to be auxotrophic for asparagine, cysteine, and threonine, while only the requirement for cysteine was experimentally observed.

### Prediction of potential metabolic drug targets

Streptococci have acquired resistance to major antibiotics which can make a successful treatment difficult (38, 39). Thus, new antibiotics are needed. As a starting point and to accelerate the discovery of new drugs metabolite essentiality analysis can be used (40). This approach removes *in-silico* each metabolite from the metabolic model. If the removal of the metabolites prevents the formation of biomass, it is deemed essential. Because this leads to a large list of metabolites with many unlikely candidates (e.g., currency metabolites such as ATP), subsequent filtering steps are needed.

To minimize potential side effects, we first removed all metabolites that also occur in the human metabolic model (41). In addition, we BLASTed the genes of all reactions that are associated with one of the essential metabolites against the human genome; if a distant homolog (e-value<0.01) for at least one gene was found, the according metabolite was also discarded from further analysis.

We applied this approach to the three reconstructed *Streptococcus* metabolic networks, identifying 10 different essential metabolites likely not involved in human metabolism (Supplemental Table S4). These drug target candidates are processed in three different pathways. The reliability of our analysis is demonstrated by the prediction of PABA (4-Aminobenzoate) as a potential drug target: this substance is already targeted by many sulfonamide antibiotics, which inhibit the dihydropteroate synthase. This enzyme is essential in bacteria for producing folate, while human acquire folate as part of their nutrition.

## Conclusions

At >95%(Table 1), the accuracy of gene knockout predictions from our *Streptococcus* models exceeds that of the most intensely curated other bacterial metabolic network, the iJO1366 model for *E. coli* (90.8%) (42). Despite this high accuracy, our model reconstructions are more conservative than those of many manually curated or automatic metabolic models: at most 4% of enzymatic reactions are not associated with a gene product (Supplemental Table 5), while the corresponding number is 6% for the iJO1366 *E. coli* model and >7% for the KBase *Streptococcus* models. Less than 2%of enzymatic reactions in our models lacked an Enzyme Commission (EC) number, whereas this is the case for >7% of reactions in the KBase models and for more than one third of enzymatic reactions in the iJO1366 model (note that the KBase models did not contain any EC numbers; we obtained these values by mapping the KBase reactions to the ModelSEED Database (8)). We conclude that the general approach demonstrated here opens up the prospect to fully automate the reconstruction of high-quality genome-scale metabolic models. The reconstruction quality could be increased even further if multiple sets of growth data are employed, *e.g.*, by including high-throughput phenotyping data (43).

## Methods

We devised a pipeline for metabolic network reconstruction suitable for full automation. This algorithm examines different sources of information in ascending order of reliability; at each step, information from the previous step is refined and overwritten.

### Base model reconstruction

We started by uploading the three genome sequences of *S. pyogenes* NZ131 (44), *S. sanguinis* SK36 (45), and *S. agalactiae* (46) to KBase (47). KBase outputs a first draft metabolic model based on sequence similarities of the genes to its database, containing reaction IDs and associated Boolean gene-protein-reaction associations (GPR rules). We removed reactions if their GPR was invalid, *i.e.*, if the reaction required at least one unknown gene to be active. The remaining reactions were carried over to the next step only if the KBase reaction abbreviation was identical to a reaction ID in the BiGG database (25).

We updated this first draft with information from the existing metabolic model of *Lactococcus lactis* (21), a close relative of Streptococci, which uses the metabolite and reaction nomenclature of the BiGG database. For each gene in the *L. lactis* genome (48), we identified homologs in the *Streptococcus* genomes by identifying reciprocal BLAST hits with e-values < 10^−13^. In some cases, this strategy resulted in the mapping of several paralogous genes to one gene annotated in the *L. lactis* genome (see below). Overall, we found *L. lactis* homologs for 62% of *S. pyogenes*, 55% of *S. sanguinis*, and 61% of *S. agalactiae* genes.

We added or updated reactions of the initial KBase model with *L. lactis* reactions if the corresponding GPR could be fulfilled with *Streptococcus* genes that had a reciprocal BLAST hit with e-value <10^−13^ between the two genomes. If a reaction in the *L. lactis* model was not associated with any gene, the corresponding (empty) GPR was also considered valid. We also included the biomass reaction of *L. lactis* into the base models. We adapted the relative abundance of the nucleotides to the G+C content observed in the *Streptococcus* genomes (*L. lactis*: G+C=35.8%, *S. sanguinis*: G+C=43.4%, *S. pyogenes*: G+C=38.5%, *S. agalactiae*: G+C=35.6%).

### Obtaining transport reactions from TransportDB

We replaced GPRs for transport reactions and added transport reactions with predictions from TransportDB (22). Transport reactions with empty GPRs that could not be filled through TransportDB were retained. Note that predicting transporters is still challenging and is an important source of inaccuracies in metabolic network reconstructions (6).

### Initial model curation using KAAS

For some reactions, the homolog prediction through bidirectional blast hits resulted in GPRs with unrealistically large paralogous gene sets. Furthermore, some reactions in the *L. lactis* template metabolic network or in the KBase predictions may have erroneous GPR associations. To reduce these two sources of error, we refined each reaction in the extended base model by comparing the GPR with the corresponding gene function predictions made by the KEGG automated annotation server (KAAS) (23). We first re-annotated all genes in the *Streptococci* genomes using KAAS, which associated metabolic genes recognized by KAAS with EC numbers and KEGG reaction identifiers. Using KEGG (49), we additionally associated each reaction in the extended base model with an EC number and a KEGG reaction identifier. We added genes to the GPR of all reactions with the same EC number as that assigned by KAAS to the gene, and we removed genes from GPRs if the gene’ s EC number according to KAAS differed from that of the reaction. If KEGG contained information on protein complexes, we introduced that information into the corresponding GPRs. We dropped reactions that lost a valid GPR association in this process.

For example, the *L. lactis* metabolic network reconstruction contains a coproporphyrinogen oxidase (CPPPGO, EC: 1.3.3.3), which requires oxygen and is associated with the *L. lactis* gene iNF518 (HemN). The KAAS annotation revealed that this gene instead encodes an oxygen-independent coproporphyrinogen-III oxidase (CPPPGO2, EC: 1.3.99.22) (50). The oxygen dependent coproporphyrinogen oxidase erroneously included in the *L. lactis* model is catalyzed by HemF, which is present neither in the genome of anaerobically living *Streptococci* nor in *L. lactis*.

### ATP maintenance reactions

The non-growth associated maintenance reaction (NGAM) accounts for ATP requirements not directly related to cell growth (*e.g.*, maintenance of turgor pressure); conversely, the growth associated maintenance reaction (GAM) accounts for energy requirements directly related to cell replication (*e.g.*, synthesis of proteins, DNA, and RNA) and is part of the biomass reaction. Appropriate rates for these ATP maintenance reactions are usually determined by growth experiments (6). Because such experiments were not available for *S. sanguinis, S. pyogenes*, or *S. agalactiae*, we set the lower bounds of the according reactions to the values appropriate for *L. lactis* (GAM: 39 mM/gDW/h; NGAM: 0.92 mM/gDW/h) (21).

### Simulated environments

The resulting draft models for *S. sanguinis* and *S. pyogenes* were used as the basis for further refinement through high-throughput gene knockout data. All analyzed gene knockout studies were performed anaerobically on undefined rich media (brain heart infusion for *S. sanguinis* (36), Todd-Hewitt Yeast medium for *S. pyogenes* (51), and TS media for *S. agalactiae* (52)). We thus allowed the uptake of all nutrients for which a transport reaction was included in the curated base model except for oxygen, constraining the lower bound of the oxygen exchange reaction to zero and the lower bound of all other exchange reactions to −5 mM/gDW/h.

### Identification of FPp and FNp

Using this set of parameters, we performed flux balance analysis (FBA) (24) with the biomass reaction provided by the *L. lactis* metabolic model. We identified FPp (false positive predictions) as those cases where our *in-silico* gene knockout simulation predicted growth (*vbiomass*>0), while the *in vivo* experiment showed no growth. Correspondingly, we identified FNp (false negative predictions) as those cases where the *in silico* gene-knockout analysis predicted no growth, while the knockout was viable in the experiment. Because no lower threshold for growth was used in the gene knockout experiments, we also interpreted any flux through the biomass reaction >10^−9^ as *in silico* growth.

### Automated network refinement with GLOBALFIT

The draft networks were refined with gene knockout libraries using the previously published GLOBALFIT algorithm (12). GLOBALFIT is a bi-level optimization program that identifies smallest sets of network modifications in order to minimize the number of FNp and FPp cases; allowed network modifications are (i) removals or (ii) reversibility changes of existing reactions; (iii) additions of reactions to the model from a database of potential reactions; (iv) removals of metabolites from the biomass; and (v) additions of metabolites to the biomass.

The strength of GLOBALFIT is its ability to consider several growth or non-growth cases simultaneously. In particular, when considering FPp, it is important to simultaneously consider a true growth case to avoid trivial solutions such as the removal of an essential reaction(12). We initially solved each FPp simultaneously with a wild type growth case and each FNp simultaneously with a non-growth case. If the network changes suggested by GLOBALFIT introduced new errors by converting true positive predictions (TPp) to FNp, or true negative predictions (TNp) to FPp, we solved the examined case again, this time simultaneously with all cases converted to false predictions by the original modification.

### Allowed model changes

For each network, we created a conservative set of potential additional reactions by blasting the corresponding genome against all genes annotated in networks contained in the BiGG database (25). As before, we identified homologous genes as bidirectional blast hits with e-values <10^−13^. All reactions with a valid GPR of homologous genes was added to the set of potential additional reactions. Streptococci are gram positive bacteria; we thus removed all reactions that do not occur in the cytosol or the extracellular space of their original metabolic model. We allowed the potential reversal of irreversible reactions only for those reactions that were classified at least as “ reversible with uncertainty” in the *E. coli* metabolic network (26).

Biomass reaction changes (addition or removal of biomass components) suggested by GLOBALFIT were manually compared to the literature and the biomass reaction generated by KBase (see Supplemental data); a fully automated approach could instead use a preformed database of potential biomass components. If a potential additional biomass component was part of the biomass reaction of the KBase model, the stoichiometric coefficient was carried over; otherwise, it was arbitrarily set to 10^−5^. No growth experiments for *Streptococci* were available to refine the coefficients. However, the exact coefficient of each biomass component is less important than its presence for gene knockout analyses (6), and corresponding inaccuracies should not significantly bias our results. However, to allow quantitative predictions of biomass production rates, the biomass stoichiometry of the final networks will require additional curation.

### Removal of exchange reactions

GLOBALFIT suggested the removal of several exchange reactions. This may indicate the absence of the corresponding transporters from the *Streptococcus* genome. However, most of these removals may simply indicate that the corresponding nutrient was missing from the undefined growth medium (12). Supplemental Table S6 lists the exchange reactions removed for each *Streptococcus* strain.

### Growth on chemically defined media for S. sanguinis

The metabolic network of *S. sanguinis* was further refined by using the gene knockout library performed on a chemically defined medium (36). In contrast to the knockout experiments on undefined media, the corresponding publication lists knockout growth rates rather than simply stating growth or non-growth. For both experiment and simulations, we considered growth rates >80% of the wildtype as growth cases, and growth rates <10% of the wildtype as non-growth cases. The metabolic network of *S. sanguinis* was then further refined using the same strategy as described above.

We could successfully predict growth of the *S. sanguinis* metabolic network on B 48 (34), SY and M3 media (35). The study of Rogers also provided growth information in media lacking one specific metabolite (riboflavin, pantheonate, thiamine, nicotinic acid, pyridoxal, folic acid, aminobenzoic acid and biotin). The *S. sanguinis* network successfully predicted the essentiality and non-essentiality of these metabolites, except the essentiality of pyridoxal and riboflavin. Therefore, we added these components to the biomass objective function. These metabolites were also part of the biomass reaction of the network reconstructed by the KBase.

The genome of *S. sanguinis* contains a gene cluster that allows the organism to produce cobalamin (vitamin B12) anaerobically; this region was presumably acquired via horizontal gene transfer (45). 10 reactions from this pathway are missing from the draft genome reconstruction. However, as cobalamin is not part of the biomass reaction and all enzymes involved in cobalamin production are non-essential, GLOBALFIT did not attempt to complete the pathway. We used GLOBALFIT to identify a minimal set of missing reactions for cobalamin production by using a growth case with a cobalamin-consuming reaction as the objective function and allowing the addition of all reactions in the BiGG database. The predicted complete pathway was identical to the one in the metabolic network of *Clostridium ljungdahlii* (53) and *Geobacter metallireducens* (54). This manual refinement did not affect the knock-out predictions.

## Acknowledgements

We acknowledge financial support through the German Research Foundation DFG (GRK 1525 supporting DH; EXC 1028 and CRC 680 to MJL).

## Author contributions

DH conceived of the study and performed all analyses except those on the two published strains, which were performed by CJF. MJL guided the study. DH and MJL wrote the manuscript. DH, CJF, and MJL edited the manuscript.

